# The interplay between host genetics and the gut microbiome reveals common and distinct microbiome features for human complex diseases

**DOI:** 10.1101/2019.12.26.888313

**Authors:** Fengzhe Xu, Yuanqing Fu, Ting-yu Sun, Zengliang Jiang, Zelei Miao, Menglei Shuai, Wanglong Gou, Chu-wen Ling, Jian Yang, Jun Wang, Yu-ming Chen, Ju-Sheng Zheng

## Abstract

There is increasing interest about the interplay between host genetics and gut microbiome on human complex diseases, with prior evidence mainly derived from animal models. In addition, the shared and distinct microbiome features among human complex diseases remain largely unclear. We performed a microbiome genome-wide association study to identify host genetic variants associated with gut microbiome in a Chinese population with 1475 participants. We then conducted bi-directional Mendelian randomization analyses to examine the potential causal associations between gut microbiome and human complex diseases. We found that *Saccharibacteria* (also known as *TM7* phylum) could potentially improve renal function by affecting renal function biomarkers (i.e., creatinine and estimated glomerular filtration rate). In contrast, atrial fibrillation, chronic kidney disease and prostate cancer, as predicted by the host genetics, had potential causal effect on gut microbiome. Further disease-microbiome feature analysis suggested that gut microbiome features revealed novel relationship among human complex diseases. These results suggest that different human complex diseases share common and distinct gut microbiome features, which may help re-shape our understanding about the disease etiology in humans.

## Introduction

Ever-increasing evidence has suggested that gut microbiome is involved in many physiological processes, such as energy harvest, immune response, and neurological function^1–3^. With successes of investigation into the clinical application of fecal transplants, modulation of gut microbiome has emerged as a potential treatment option for some complex diseases, including inflammatory bowel disease and colorectal cancer ^4, 5^. However, it is still unclear whether the gut microbiome has the potential to be clinically applied for the prevention or treatment of many other complex diseases. Therefore, it is important to clarify the bi-directional causal association between gut microbiome and human complex diseases or traits.

Mendelian randomization (MR) is a method that uses genetic variants as instrumental variables to investigate the causality between an exposure and outcome in observational studies^6^. Prior literature provides evidence that the composition or structure of the gut microbiome can be influenced by the host genetics^7–10^. On the other hand, host genetic variants associated with gut microbiome were rarely explored in Asian populations, thus we are still lacking instrument variables to perform MR for gut microbiome in Asians. This calls for novel microbiome genome-wide association study (GWAS) in Asian populations.

Along with the causality issue between the gut microbiome and human complex diseases, it is so far unclear whether human complex diseases had similar or unique gut microbiome features. Identifying common and distinct gut microbiome features across different diseases might shed light on novel relationships among the complex diseases and update our understanding about the disease etiology in humans. However, the composition and structure of gut microbiome are influenced by a variety of factors including environment, diet and regional variation ^11–13^, which posed a key challenge for the description of representative microbiome features for a specific disease.

Although there were several studies comparing disease-related gut microbiome ^14–16^, few of them has examined and compared the microbiome features across different human complex diseases.

In the present study, we performed a microbiome GWAS in a Chinese cohort study: the Guangzhou Nutrition and Health Study (GNHS)^17^, including 1475 participants. Subsequently, we applied a bi-directional MR method to explore the genetically predicted relationship between gut microbiome and human complex diseases. To explore novel relationships among human complex diseases based on gut microbiome, we investigated the shared and distinct gut microbiome features across diverse human complex diseases ^18^.

## Result

### Overview of the study

Our study was based on the GNHS, with 4048 participants (40-75 years old) living in urban Guangzhou city recruited during 2008 and 2013 ^17^. In the GNHS, stool samples were collected among 1937 participants during follow-up visits, among which 1475 unrelated participants without taking anti-biotics were included in our discovery microbiome GWAS. We then included additional 199 participants with both genetic and gut microbiome data as a replication cohort, which belonged to the control arm of a case-control study of hip fracture in Guangdong Province, China ^19^.

For both discovery and replication cohorts, genotyping was carried out with Illumina ASA-750K arrays. Quality control and relatedness filters were performed by PLINK1.9 ^20^. We conducted the genome-wide genotype imputation with 1000 Genomes Phase3 v5 reference panel by Minimac3^21–23^. HLA region was imputed with Pan-Asian reference panel and SNP2HLA v1.0.3 ^24–26^.

### Association of host genetics with gut microbiome features

We performed a series of microbiome GWAS with PLINK 1.9 based on logistic models for binary variables ^20^. For continuous variables, we used GCTA with mixed linear model-based association (MLMA) method ^27, 28^. We also analyzed categorical variable enterotypes of the participants based on genus-level relative abundance of gut microbiome, using the Jensen-Shannon Distance (JSD) and the Partitioning Around Medoids (PAM) clustering algorithm ^29^. The participants were subsequently clustered into two groups according to the enterotypes (*Prevotella* vs *Bacteroides*). Thereafter, we performed GWAS for enterotypes using logistic regression model to explore potential associations between host genetics and enterotypes. However, we did not find any genome-wide significant locus with p<5×10^-8^. Furthermore, we used a restricted maximum likelihood analysis (REML) with GCTA to estimate the SNP-based heritability, and the estimate heritability of the enterotype was 0.055 (SE=0.19, Supplementary Table S2) ^30^.

To examine the association of host genetic variants with alpha diversity, we performed GWAS for three indices (Shannon diversity index, Chao1 diversity indices and observed OTUs index), but again no genome-wide significant signal (p<5×10^-8^) was found. In the discovery cohort, the heritability of alpha diversity ranged from 0.054 to 0.14 (SE=0.20 for all indices, Supplementary Table S2). To further investigate if there is host genetic basis underlying alpha diversity, we constructed a polygenic score for each alpha diversity indicator in the replication cohort, using the genetic variants which showed suggestive significance (p<5×10^-5^) in the discovery GWAS. The polygenic score was not significantly associated with its corresponding alpha diversity index in our replication cohort. Meanwhile, none of the associations with alpha diversity indices reported in the literature could be replicated (Supplementary Table S7) ^7^.

We performed a beta diversity GWAS using a tool called MicrobiomeGWAS ^31^, and found that one locus at *SMARCA2* gene (rs6475456) was associated with beta-diversity at a genome-wide significance level (p=3.96×10^-9^). However, we could not replicate the results in the replication cohort, which may be due to the limited sample size of the replication cohort. In addition, prior literature had reported 73 genetic variants that were associated with beta diversity ^8, 13, 32, 33^, among which we found that 3 single nucleotide polymorphisms (SNP, *UHRF2* gene-rs563779, *LHFPL3* gene-rs12705241, *CTD-2135J3*.*4*-rs11986935) had nominal significant (p<0.05) association with beta-diversity in our cohort (Supplementary Table S6), although none of the association survived Bonferroni correction. These studies used various methods for the sequencing and calculation of beta diversity, which raised challenges to verify and extrapolate results across populations.

We then took the genetic loci reported to be associated with individual taxa in prior studies ^7, 8, 13, 33^ for replication in our GNHS dataset. Although there are still some signals with nominal significance (p<0.05) in our study (e.g., 7 loci associated with *Lachnospiraceae*, *Coprococcus* or *Bacteroides* with p<0.05; Supplementary Table S5), none of the associations of these genetic variants with taxa survived the Bonferroni correction (p<1×10^-4^). The null results may be because of various clustering similarities, classifiers or reference databases to annotate taxa and different sequencing methods used in these studies.

We subsequently performed GWAS discovery for individual gut microbes in our own GNHS discovery dataset. For the taxa present at more than ninety percent of participants and alpha diversity, we used Z-score normalization to transform the distribution and carried out analysis based on a log-normal model. A MLMA test in the GCTA was used to assess the association, with the first five principal components, age, sex and sequencing batch fitted as fixed effects and the effects of all the SNPs fitted as random effects ^27, 28, 34^. For other taxa present at fewer than ninety percent, we transformed the absence/presence of the taxon into binary variables and used PLINK1.9 to perform a logistic model, adjusted for the first five principal components, age, sex and sequencing batch.

For all the gut microbiome taxa, the significant threshold was defined as 5×10^-8^ in the discovery stage. As some taxa were correlated with each other, we also used an eigendecomposition analysis to calculate the effective number of independent taxa on each taxonomy level (phylum level: 2.3, class level: 2.9, order level: 2.9, family level: 5.5, genus level: 5.6, species level: 3.2) ^35, 36^. We found that 6 taxa were associated with host genetic variants in the discovery cohort (p<5×10^-8^/n, n is the effective number of independent taxa on each taxonomy level, Supplementary Table S4); however, these associations were not significant (p>0.05) in the replication cohort. We then used a threshold of p<5×10^-5^ at the GWAS discovery stage to incorporate additional genetic variants which may explain a larger proportion of heritability for taxa, based on which we constructed a polygenic score for each taxon in the replication. We found that the polygenic scores were significantly associated with 5 taxa including *Saccharibacteria* (also known as *TM7* phylum), *Clostridiaceae*, *Comamonadaceae*, *Klebsiella* and *D168* species (belong to *Desulfovibrio* genus) in the replication set (p<0.05, Methods, see also Figure 1A, 1B, 1C, 1D, 1E).

**Figure 1.**
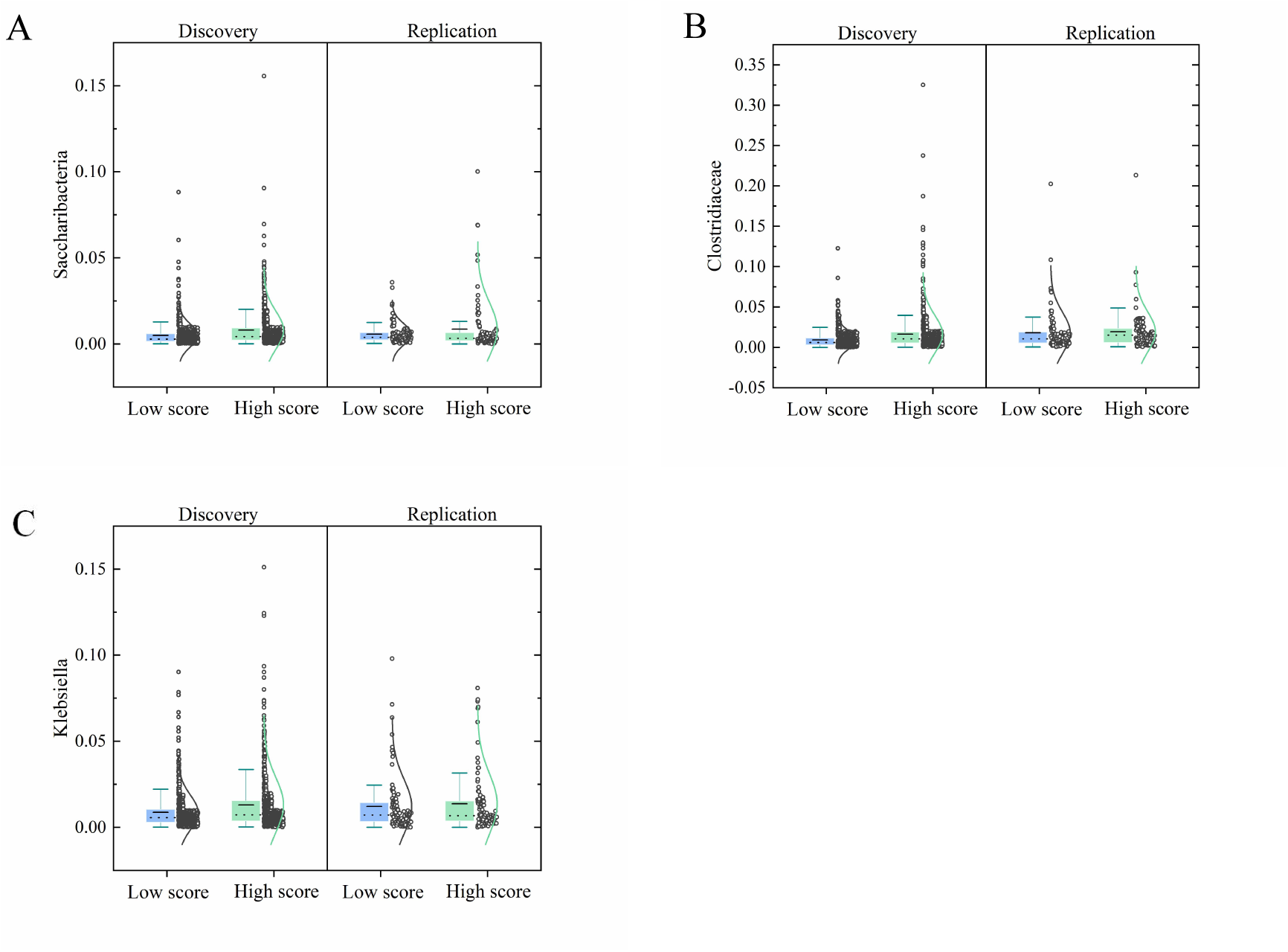
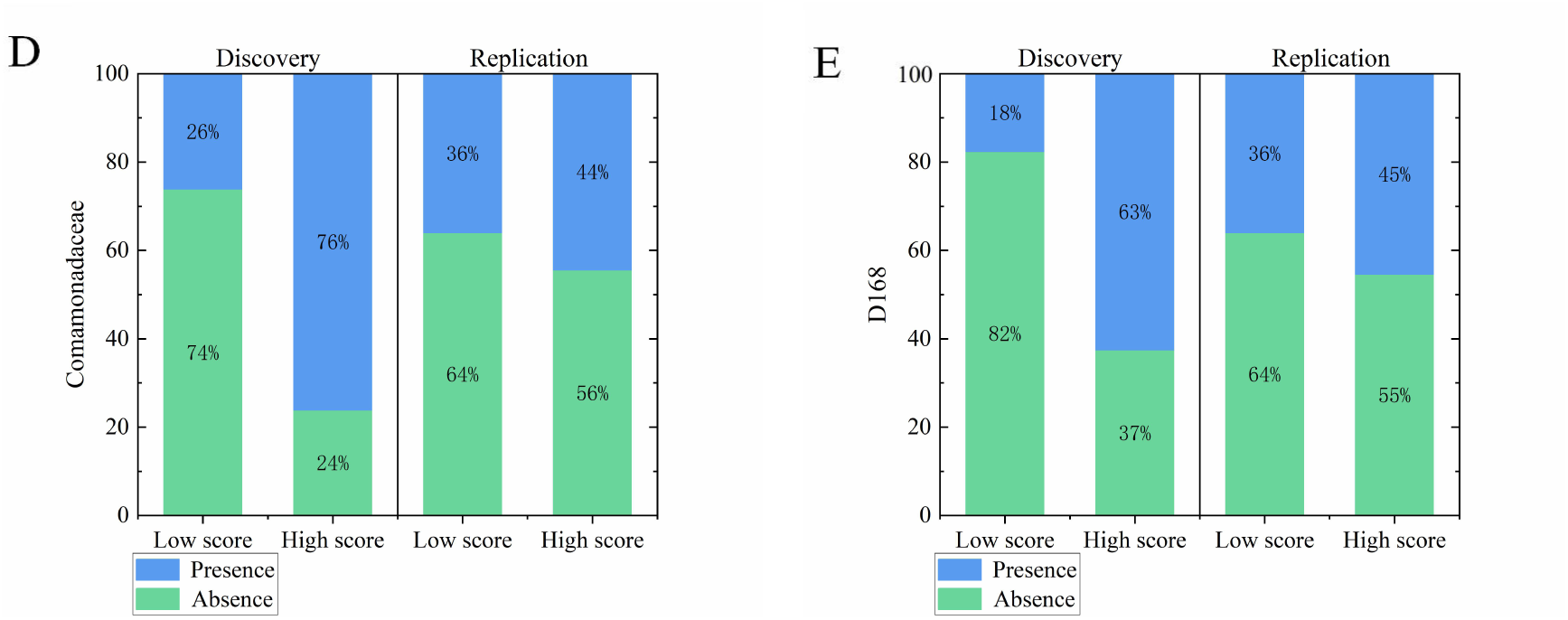
Association of host polygenic score with gut microbiome. The participants were divided into high and low polygenic score group according to median levels of the polygenic score. The dots on the right of the box represent the distribution of polygenic score. The dash line in the box is the position of median line and the solid line is the position of mean line. The length of box depends on upper quartile and lower quartile of datum. Sample size at the discovery stage is 1475, and that at replication stage is 199. **(A).** Correlation of *Saccharibacteria* abundance with the polygenic score (including 51 lead SNPs, Supplementary Table S8). **(B).** Correlation of *Clostridiaceae* abundance with the polygenic score (including 37 lead SNPs, Supplementary Table S8). **(C).** Correlation of *Klebsiella* abundance with the polygenic score (including 47 lead SNPs, Supplementary Table S8). **(D).** Correlation of *Comamonadaceae* presence with the polygenic score (including 37 lead SNPs, Supplementary Table S8). **(E).** Correlation of *D168* species presence with the polygenic score (including 34 lead SNPs, Supplementary Table S8).

### Genetic correlation of gut microbiome and traits

We used GCTA to perform a bivariate GREML (genomic-relatedness-based restricted maximum-likelihood) analysis to estimate the genetic correlation between gut microbiome and traits in the GNHS ^27, 37^. The traits included BMI, fasting blood sugar (FBS), glycosylated hemoglobin (HbA1c), systolic blood pressure (SBP), diastolic blood pressure (DBP), high density lipoprotein cholesterol (HDL-C), low density lipoprotein cholesterol (LDL-C), total cholesterol (TC) and triglyceride (TG), none of which could pass Bonferroni correction. Additionally, HDL-C was the only trait that had nominal genetic correlation (p<0.05) with gut microbes (specifically, *Desulfovibrionaceae* and *[Prevotella]*, Supplementary Table S3).

### Bi-directional assessment of the genetically predicted association between gut microbiome and complex diseases/traits

Using genetic variants-composed polygenic scores as genetic instruments, we performed MR analysis to assess the putative causal effect of microbiome (*Saccharibacteria*, *Clostridiaceae*, *Comamonadaceae*, *Klebsiella* and *D168* species) on human complex diseases or traits. Inverse variance weighted (IVW) method was used for the MR analysis, while other three methods (Weighted median, MR-Egger and MR-PRESSO) ^38, 39^ were applied to confirm the robustness of results. The horizontal pleiotropy was assessed using MR-PRESSO Global test and MR-Egger Regression. For the analysis of gut microbiome on complex traits, we downloaded public available GWAS summary statistics of complex traits (n=58) and diseases (type 2 diabetes mellitus (T2DM), atrial fibrillation (AF), colorectal cancer (CRC) and prostatic cancer (PCa)) reported by BioBank Japan ^40–44^. The result suggested that *Saccharibacteria* could potentially decrease the concentration of serum creatinine (p=0.008, Supplementary Table S9) and increase estimated glomerular filtration rate(eGFR)(p=0.003, Supplementary Table S9), which helps improve renal function. We did not find evidence of pleiotropic effect: genetic variants associated with *Saccharibacteria* were not associated with any of the above traits (58 complex traits and 4 disease outcomes, p<0.05/62). These taxa were not causally associated with other human complex diseases or traits in our MR analyses, which may be due to the limited genetic instruments discovered in our present study.

We subsequently performed a reserve MR analysis to assess the potential causal effect of human complex diseases on gut microbiome features. For the reserve MR analyses, the diseases of interests included T2DM, AF, coronary artery disease (CAD), chronic kidney disease (CKD), Alzheimer’s disease (AD), CRC and PCa, and their instrumental variables for the MR analysis were based on the previous large-scale GWAS in East Asians^40, 45–50^. The results suggested that AF and CKD were causally associated with gut microbiome (See also Figure 2A, 2B, Supplementary Table S10). Genetically predicted higher risk of AF was associated with lower abundance of *Coprophilus*, *Lachnobacterium*, *Barnesiellaceae*, *Veillonellaceae* and *Mitsuokella*, and higher abundance of *Alcaligenaceae*. Additionally, genetically predicted higher risk of CKD could increase *Anaerostipes* abundance and higher risk of PCa could decrease *[Prevotella]*.

**Figure 2.**
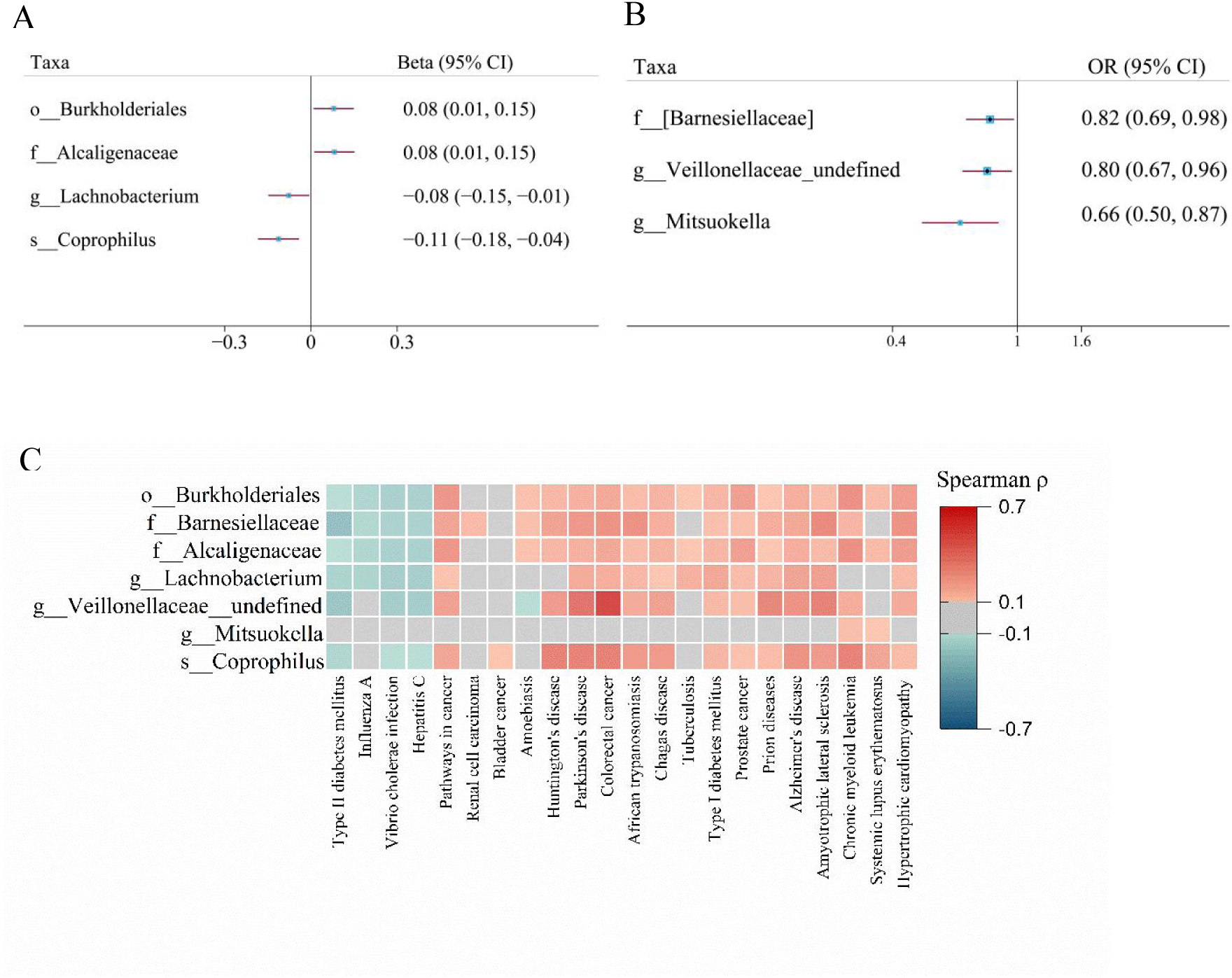
Effect of host genetically predicted higher atrial fibrillation risk on gut microbiome. **(A)**. Causal association of atrial fibrillation with abundance of *Burkholderiales*, *Alcaligenaceae*, *Lachnobacterium* and *Coprophilus*. The effect sizes of atrial fibrillation on taxa are changes in abundance of bacteria (10-SD of log-transformed) per genetically determined higher log odds of atrial fibrillation. **(B)**. Causal association of atrial fibrillation with presence of *Barnesiellaceae*, *Veillonellaceae_undefined* and *Mitsuokella*. The effect size of atrial fibrillation on taxa are present as odds ratio increase in log odds of atrial fibrillation. **(C)**. The heat map shows correlation of AF-associated taxa with predicted diseases. The grey components show no significance of correlation with Bonferroni correction (p>0.05/ (5.6*22), p>0.0004).

To further investigate the potential complex diseases that may be correlated with the taxa affected by AF, we applied Phylogenetic Investigation of Communities by Reconstruction of Unobserved States (PICRUSt) to predict the disease pathway abundance ^51^. We used Spearman’s rank-order correlation to test whether 22 human complex diseases were associated with the aforementioned AF-associated taxa (See also Figure 2C). The heatmap indicated that cancers and neurodegenerative diseases including Parkinson’s disease (PD), AD, amyotrophic lateral sclerosis (ALS) as well as AF were correlated with similar gut microbiome. Although the association among these diseases are highly supported by previous studies ^52–54^, no study has compared common gut microbiome features across these different diseases.

### Microbiome features of human complex diseases

To compare gut microbiome features across human diseases, we chose 22 human complex diseases from predicted abundance and performed k-medoids clustering ^18^. We used an *m × n* matrix to perform the cluster analysis, where m is the number of participants and n is the number of selected diseases. According to optimum average silhouette width ^55^, we chose optimal number of clusters for further analysis (See also Figure 3A). The plot showed that neurological diseases including ALS and AD belonged to the same cluster, while PD and CRC had much similarity in gut microbiome. The results also suggested that systemic lupus erythematosus (SLE) and chronic myeloid leukemia (CML) shared similar gut microbiome features. Moreover, we could replicate these clusters in our replication cohort, which suggested that the clustering results were robust (See also Figure 3B).

**Figure 3.**
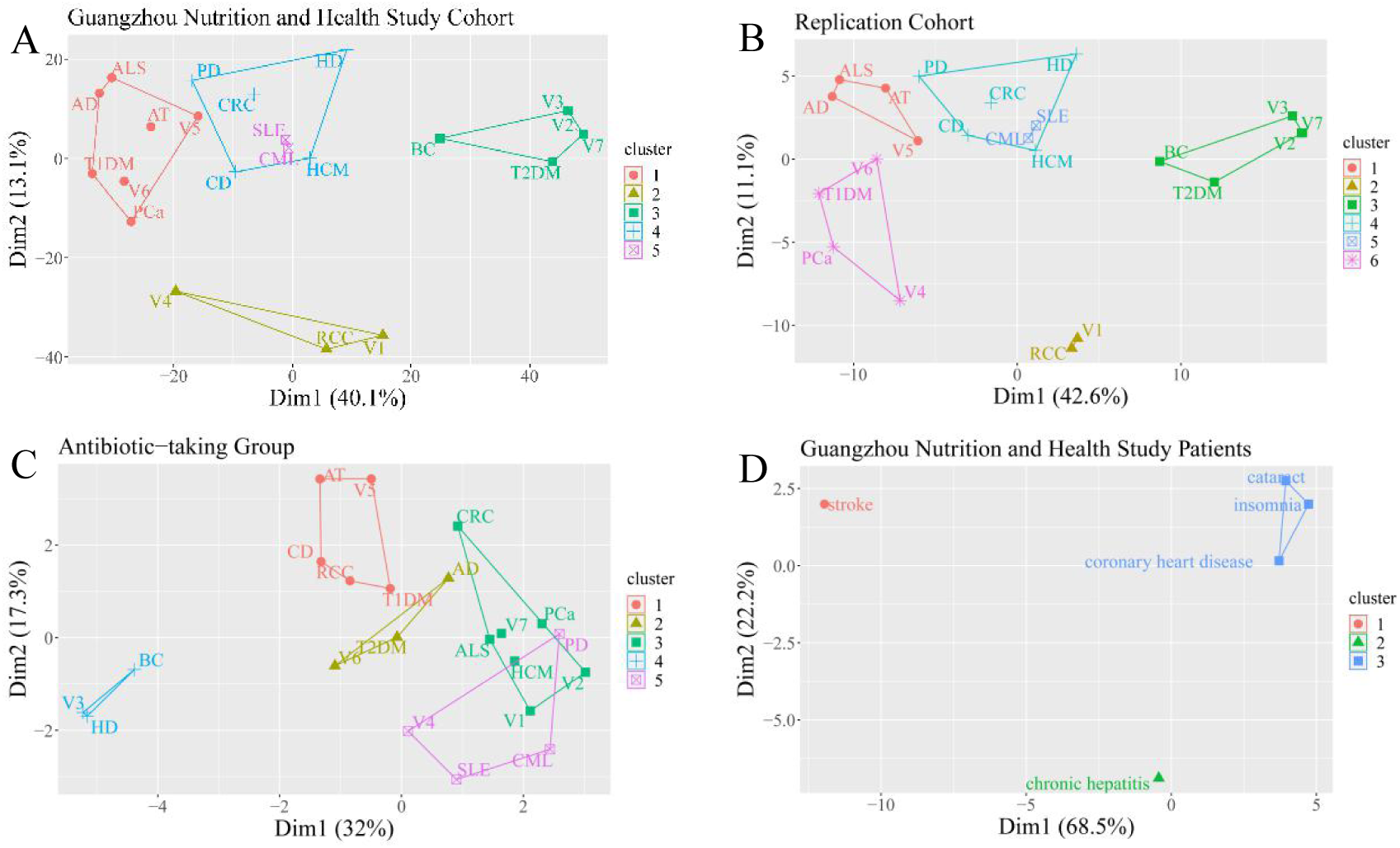
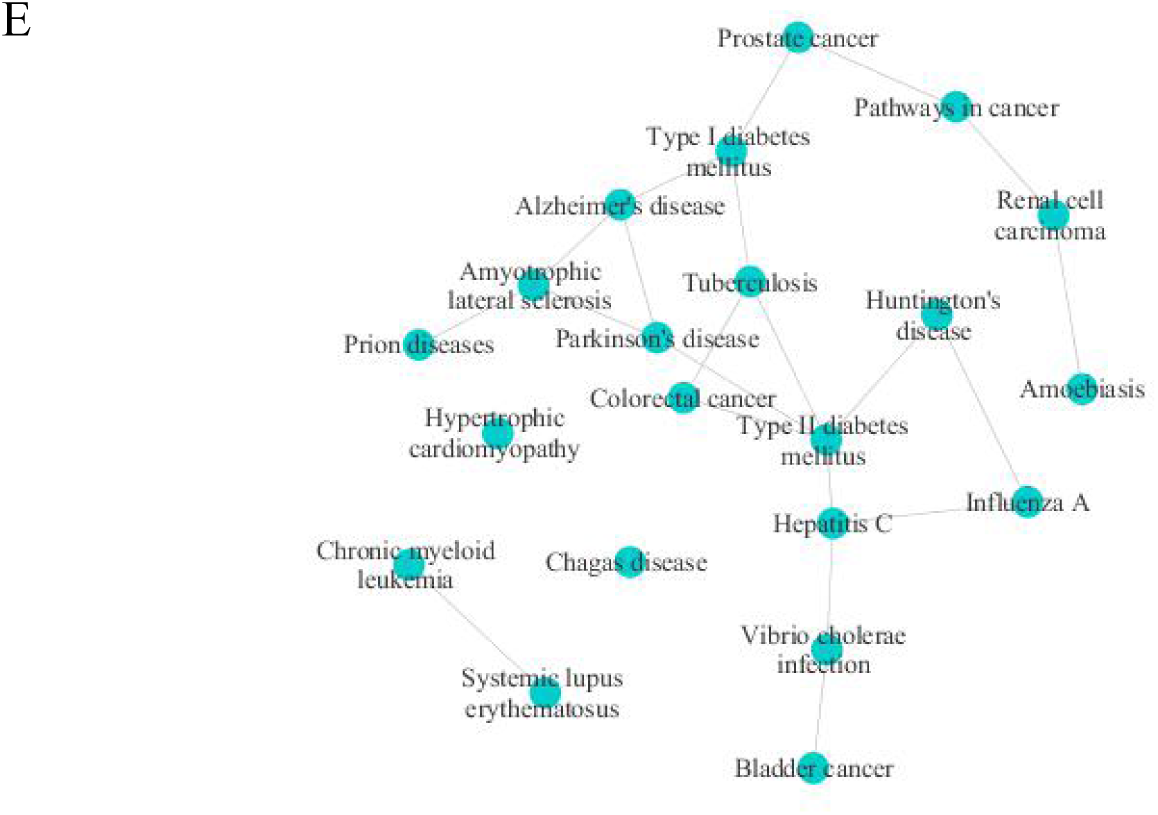
Association and cluster of diseases predicted by the gut microbiome. **(A)**. Plot of clusters in Guangzhou Nutrition and Health Study (GNHS) cohort (n=1919). (**B)**. Plot of cluster results in the replication cohort (n=217). **(C)**. Plot of 5 clusters in antibiotic-taking participants (n=18). The optimal cluster is 5 in GNHS cohort and 6 in the replication. The clusters share consistent components between two studies. In contrast, components are different between antibiotic-taking participants and control groups. Dimension1 (Dim1) and dimension2 (Dim2) can explain 40.1% and 13.1% variance, respectively in GNHS cohort. The annotation for variables is as following. AT: African trypanosomiasis, AD: Alzheimer’s disease, V1: Amoebiasis, ALS: Amyotrophic lateral sclerosis, BC: Bladder cancer, CD: Chagas disease, CML: Chronic myeloid leukemia, CRC: Colorectal cancer, V2: Hepatitis C, HD: Huntington’s disease, HCM: Hypertrophic cardiomyopathy, V3: Influenza A, PD: Parkinson’s disease, V4: Pathways in cancer, V5: Prion disease, PCa: Prostate cancer, RCC: Renal cell carcinoma, SLE: Systemic lupus erythematosus, V6: Tuberculosis, T1DM: Type I diabetes mellitus, T2DM: Type II diabetes mellitus, V7: Vibrio cholerae infection. **(D)**. Plot of clusters in GNHS patients. Patients get only one of the follow diseases: stroke (n=8), chronic hepatitis (n=19), coronary heart diseases (n=40), cataract (n=124) and insomnia (n=68). **(E)**. Gut microbiome-predicted network of relationship among different human complex diseases. The interaction is determined by SPIEC-EASI with non-normalized predicted abundance data.

We further asked whether gut microbiome contributed to the novel clustering. To this end, we repeated the analysis among participants who took antibiotic less than two weeks before stool sample collection, considering that antibiotic treatments were believed to cause microbiome imbalance, and the clusters were totally different in this group (See also Figure 3C). The results indicated a totally different clustering, which suggested that gut microbiome indeed contributed to the correlations among diseases. To further demonstrate common microbiome features across different diseases, we examined the correlation of the predicted diseases with genus-level taxa. The results showed that human complex diseases had shared similar gut microbiome features, as well as distinct features on their own (See also Figure 4).

**Figure 4.**
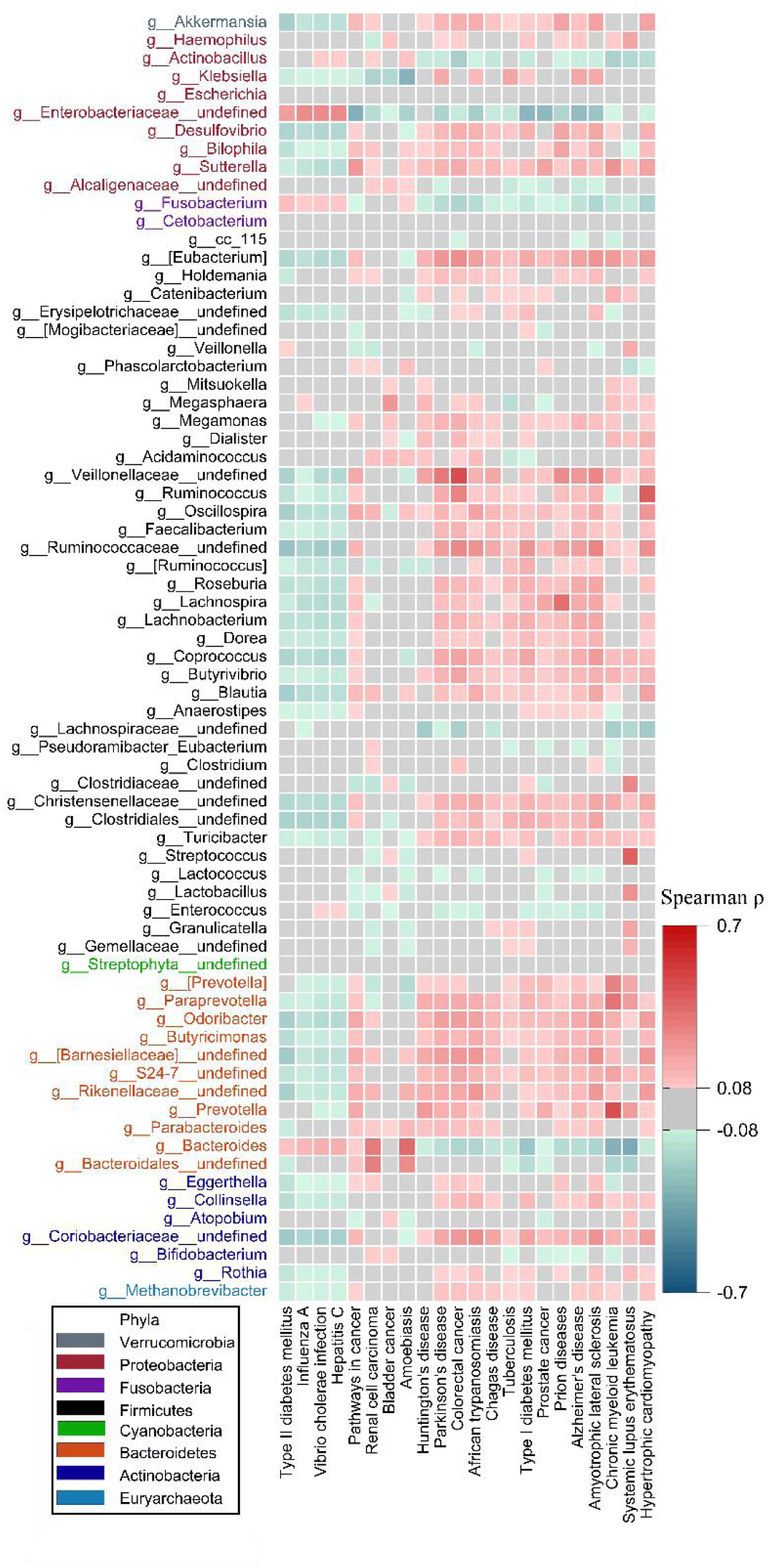
Correlation of the human complex diseases with gut microbiome on genus level. The heat map shows correlation of predicted diseases and gut microbiome on genus level. The grey components show no significance of correlation with Bonferroni correction (p>0.05/ (5.6*22), p>0.0004).

To validate the accuracy of the association between the predicted disease-related gut microbiome features and the corresponding disease, we used T2DM as an example, examining the association of predicted T2DM-related microbiome features with T2DM risk in our GNHS samples. We constructed a microbiome risk score (MRS) based on 16 selected taxa with predicted correlation coefficient with T2DM greater than 0.2. A logistic regression was used to examine the above MRS with T2DM risk in the GNHS (n=1886, with 217 T2DM cases). The results showed that higher MRS was associated with lower risk of T2DM (odds ratio: 0.850, 95% confidence interval: 0.804 to 0.898, p=8.75×10^-9^).

Based on the above results, we proposed a hypothesis that related diseases might share similar gut microbiome features. To test for this hypothesis, we performed validation analysis by including GNHS participants who had one of the following self-reported diseases: stroke (n=8), chronic hepatitis (n=19), coronary heart diseases (CHD) (n=40), cataract (n=124) and insomnia (n=68). The results of k-medoids clustering suggested that CHD, cataract and insomnia shared common gut microbiome features, which was supported by the prior research reporting that both patients suffering insomnia and women receiving cataract extraction had increased risks of CHD ^56–58^.

## Discussion

Our study is among the first to investigate the host genetics-gut microbiome association in East Asian populations and reveals that several microbiome species (e.g., *Saccharibacteria* and *Klebsiella*) are influenced by host genetics. We then show that *Saccharibacteria* might causally improve renal function by affecting renal function biomarkers (i.e., creatinine and eGFR). On the other hand, complex diseases such as atrial fibrillation, chronic kidney disease and prostate cancer, have potential causal effect on gut microbiome. More interestingly, our results indicate that different human complex diseases may be mechanically correlated by sharing common gut microbiome features, but also maintaining their own distinct microbiome features.

Previous studies and our study showed that gut microbiome had an inclination to be influenced by host genetics ^8, 10, 33, 59^, although the successful replication tends to be rare. We could not validate any of the reported genetic variants that were significantly associated with gut microbiome features in prior reports, which may reflect the difference in population and heterogeneity between study but also raise concerns about the reproducibility. Many factors including ethnic differences, gene-environment interaction and dissimilarity in sequencing methods may make it hard to extrapolate results from microbiome GWAS across populations in the microbiome field. Nevertheless, we successfully replicate several polygenic scores of gut microbiome, and the current study represent the largest dataset in Asian populations and would be a unique resource to be used in large-scale trans-ethnic meta-analysis of microbiome GWAS in future.

The MR analysis of gut microbiome on diseases or traits found that *Saccharibacteria* might decrease the concentration of serum creatinine and increase eGFR. Little is known about the *Saccharibacteria* as one of the uncultivated phyla, and it might be essential for the immune response, oral inflammation and inflammatory bowel disease as shown previously^60–62^. Our results also provide genetic instrument of *Saccharibacteria* for further causal analysis with other complex diseases. The reverse MR analysis provided evidence that AF, CKD and PCa could causally influence gut microbiome. As our study is among the first to investigate gene-microbiome association in East Asians, we need further study in this region to confirm our results. Additionally, rare and low-frequency variants may have an important impact on common diseases^63^, thus it will be of interest to clarify the effects of low-frequency variants on gut microbiome in cohorts with large sample sizes in future.

Our results indicate that gut microbiome helps reveal novel and interesting relationships among human complex diseases, suggesting that different diseases may have common and distinct gut microbiome features. A prior study including participants from different countries identified three microbiome clusters^29^. Notably, this study focused on classifying the individuals into distinct enterotypes regardless of the individuals’ health status, while in the present study we described representative microbiome features for diseases of interest. We provide an approach to interpret the data from mechanistic studies based on microbiome. The microbiome features revealed a close association of AF with neurodegenerative diseases as well as cancers, which was supported by prior studies showing that AF had correlation with AD and PD^52, 53^, and AF patients had relatively higher risks of several cancers including lung cancer and CRC^54, 64^. We also observed that microbiome features of SLE and CML were highly similar. Interestingly, a tyrosine kinase inhibitor of platelet-derived growth factor receptor, imatinib, was widely used to treat CML and significantly ameliorated survival in murine lupus autoimmune disease^65^. In addition, association between CRC and PD has been reported in several observational cohorts^66, 67^. Collectively, these findings strongly support our hypothesis that human complex diseases sharing similar microbiome features might be mechanically correlated. Furthermore, from the perspectives of risk genes of AF and neurodegenerative diseases, previous GWAS for AF have identified two loci at *PITX2* gene*-*rs6843082 and *C9orf3* gene*-*rs7026071, which were also associated with the risk of ALS (p=0.0138 and p=0.049, respectively) ^68–70^.

In summary, we perform bi-directional MR analyses and reveal some causal relationships between abundance of gut microbiome and human complex diseases or traits. The disease and gut microbiome feature analysis from population studies reveals novel relationships among human complex diseases, which may help re-shape our understanding about the disease etiology, as well as extending clinical indications of existing drugs for different diseases.

## Method

### Study participants and sample collection

Our study was based on the Guangzhou Nutrition and Health Study (GNHS), with 4048 participants (40-75 years old) living in urban Guangzhou city recruited during 2008 and 2013 ^17^. We followed up participants every three years. In the GNHS, stool samples were collected among 1937 participants during follow-up visits. Among those with stool samples, 1717 participants had genetic data and 1475 participants with identical by decent (IBD) less than 0.185.

We included 199 participants with both genetic and gut microbiome data as a replication cohort, which belonged to the control arm of a case-control study of hip fracture with the participants (52-83 years old) recruited between June 2009 and August 2015 in Guangdong Province, China ^19^.

Blood samples of all participants were collected after an overnight fasting and buffy coat was separated from whole blood and stored at -80℃. Stool samples were collected during the on-site visit of the participants at Sun Yat-sen University. All samples were manually stirred, separated into tubes and stored at -80℃ within four hours.

### Genotyping data

For both discovery and replicattion cohorts, DNA was extracted from leukocyte using the TIANamp® Blood DNA Kit as per the manufacturer’s instruction. DNA concentrations were determined using the Qubit quantification system (Thermo Scientific, Wilmington, DE, US). Extracted DNA was stored at -80°C. Genotyping was carried out with Illumina ASA-750K arrays. Quality control and relatedness filters were performed by PLINK1.9 ^20^. Individuals with high or low proportion of heterozygous genotypes (outliers defined as 3 standard deviation) were excluded^71^. Individuals who had different ancestries (the first two principal components ±5 standard deviation from the mean) or related individuals (IBD>0.185) were excluded^71^. Variants were mapping to the 1000 Genomes Phase3 v5 by SHAPEIT ^23, 72^ and then we conducted the genome-wide genotype imputation with 1000 Genomes Phase3 v5 reference panel by Minimac3^21, 22^. Genetic variants with imputation accuracy RSQR > 0.3 and MAF > 0.05 were included in our analysis. We used Pan-Asian reference panel consist of 502 participants and SNP2HLA v1.0.3 to impute HLA region ^24–26^.

### Sequencing and processing of 16S rRNA data

Microbial DNA was extracted from fecal samples using the QIAamp® DNA Stool Mini Kit per the manufacturer’s instruction. DNA concentrations were determined using the Qubit quantification system. The V3-V4 region of the 16S rRNA gene was amplified from genomic DNA using primers 341F and 805R. Sequencing was performed using MiSeq Reagent Kits v2 on the Illuimina MiSeq System.

Fastq-files were demultiplexed by the MiSeq Controller Software. Ultra-fast sequence analysis (USEARCH) was performed to trim the sequence for amplification primers, diversity spacers, sequencing adapters, merge-paired and quality filter^73^. Operational taxonomic units (OTUs) were clustered based on 97% similarity using UPARSE^74^.

OTUs were annotated with Greengenes 13_8 (https://greengenes.secondgenome.com/). After randomly selecting 10000 reads for each sample, Quantitative Insights into Microbial Ecology (QIIME) software version 1.9.0 was used to calculate alpha diversity (Shannon diversity index, Chao1 diversity indices and observed OTUs index) based on the rarefied OTU counts^75^.

## Statistical analysis

### Genome-wide association analysis of gut microbiome features

For each of the GNHS participants and the replication cohort, we clustered participants based on genus-level relative abundance, estimating the JSD distance and PAM clustering algorithm, and then defined two enterotypes according to Calinski-Harabasz Index^29, 76^. PLINK 1.9 was used to perform a logistic regression model for enterotypes and taxa present at fewer than ninety percent, adjusted for age, sex and the first five principal components.

For beta diversity, the analyses for the genome-wide host genetic variants with beta diversity was performed using MicrobiomeGWAS^31^, adjusted for covariates including the first five principal components, age and sex. We filtered OTUs present at fewer than ten percent of participants to calculate Bray–Curtis dissimilarity.

Alpha diversity was calculated after randomly sampling 10000 reads per sample. For the taxa present at more than ninety percent of participants and alpha diversity, we used Z-score normalization to transform the distribution and carried out analysis based on a log-normal model. A mixed linear model based association (MLMA) test in GCTA was used to assess the association, fitting the first five principal components, age, sex and sequencing batch as fixed effects and the effects of all the SNPs as random effects ^27, 28, 34^. For other taxa present at fewer than ninety percent, we transformed the absence/presence of the taxon into binary variables and used PLINK1.9 to perform a logistic model, adjusted for the first five principal components, age, sex and sequencing batch. For all the gut microbiome features, the significant threshold was defined as 5×10^-8^ /n (n is the effective number of independent taxa on each taxonomy level) in the discovery stage. We estimated genomic inflation factors with LDSC v1.0.1 at local server ^77^.

### Proportion of variance explained by all SNPs

We used the GREML method in GCTA to estimate the proportion of variance explained by all SNPs ^30^. The taxa were divided into two groups based on whether the taxa were present in the ninety percent of participants or not. For alpha diversity and taxa, our model was adjusted for constrain covariates including sex and sequencing batch, as well as quantitative covariates including the first five principal components and age. The model was adjusted for the same covariates except for sequencing batch for analysis of enterotype.

### Genetic correlation of gut microbiome and traits

We used GCTA to perform a bivariate GREML analysis to estimate the genetic correlation between gut microbiome and traits in the GNHS^27, 37^. The gut microbiome was divided into two groups according to the previous description. We used continuous variables to taxa present at more than ninety percent of participants and traits. For taxa present at fewer than ninety percent of participants, we used binary variables according to the absence/presence of taxa. This analysis included traits such as BMI, FBS, HbA1c, SBP, DBP, HDL-C, LDL-C, TC and TG.

### Constructing polygenic scores for taxa and alpha diversity

We selected lead SNPs using PLINK v1.9 with the ‘—clump’ command to clump SNPs that *p* value < 5×10^-5^ and r^2^<0.1 within 0.1 cM. We used beta coefficients as weight to construct polygenic scores for taxa and alpha diversity. For alpha diversity and taxa present at more than ninety percent participants, we constructed weighted polygenic scores and performed the analysis on a general linear model with a negative binomial distribution to test for association between the polygenic scores and taxa, adjusted for the first five principal components, age, sex and sequencing batch. We used weighted polygenic scores and logistic regression to the absence/presence taxa, adjusted for the same covariates as in the above analysis. Taxa with significance (p<0.05) in the replication cohort were included for the further analysis.

### The effective number of independent taxa

As some taxa were correlated with each other, we used an eigendecomposition analysis to calculate the effective number of independent taxa on each taxonomy level ^35, 36^. Matrix M is an m×n matrix, where m is the number of participants and n is the number of taxa on the corresponding taxa level. Matrix A is the variance-covariance matrix of matrix M. P is the matrix of eigenvectors. diag{*λ*_1_, *λ*_2_, …, *λ*_*n*_} is the diagonal matrix comprised of the ordered eigenvalues, which can be calculated as:

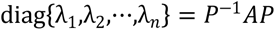

The effective number of independent taxa can be calculated as:

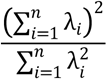

### Bi-directional MR analysis

In the analysis of potential causal effect of gut microbiome features on diseases, we used independent genetic variants (selected as part of the polygenic score analysis) as the instrument variables. For each trait, we excluded instrument variables that showed significant association with the trait (p<0.05/n, n is number of independent genetic variants). In the analysis of potential causal effect of diseases on gut microbiome features, we selected genetic variants that were replicated in East Asian populations as instrument variables. As all instrument variables were from East Asian populations, we chose independent genetic variants (r2<0.1) based on GNHS cohort. We identified the best proxy (r2 > 0.8) based on GNHS cohort or discarded the variant if no proxy was available. We used inverse variance weighted (IVW) method to estimate effect size. To confirm the robustness of results, we performed other three MR methods including weighted median, MR-Egger and MR-PRESSO. To assess the presence of horizontal pleiotropy, we performed MR-PRESSO Global test and MR-Egger Regression. Effect sizes of gut microbiome on traits were dependent on units of traits^43^ (Supplementary table S1). Results of human complex diseases on the absence/presence gut microbiome were presented as risk of presence (vs absence) of the microbiome per log odds difference of the disease. Results of diseases on other gut microbiome and alpha diversity were presented as changes in abundance of taxa (10-SD of log transformed) per log odds difference of the respective disease.

The statistical significance of gut microbiome on traits and diseases was defined as p<0.0008 (0.05/62). In addition, the statistical significance of diseases on gut microbiome features was defined as p<0.05/n (n is the effective number of independent taxa on the corresponding taxonomy level). Results that could not pass Bonferroni adjustment but p<0.05 in all four MR methods were considered as potential causal relationships. We performed MR analyses on R v3.5.3.

### Pathway analysis

We used OTUs by QIIME and annotated the variation of functional genes with Phylogenetic Investigation of Communities by Reconstruction of Unobserved States (PICRUSt) ^51^. The pathways and diseases were annotated using KEGG ^78–80^. We used Spearman’s rank-order correlation to investigate association of predicted pathway or diseases abundance with taxa. In the heatmap, diseases were clustered with ‘hcluster’ function on R. To test whether non-normalized pathway or disease abundance was associated with each other, we used SPIEC-EASI to test the interaction relationship, and then used Cytoscape v3.7.2 to visualize the interaction network ^81, 82^.

### Construction of the microbiome risk score

To construct a microbiome risk score for T2DM, we used a Spearman’s rank-order correlation to select taxa with the absolute value of correlation coefficient higher than 0.2. Score for each taxon abundance <5% quantile in our study was defined as 0. For those above 5%, score for each taxon showing negative association with T2DM was defined as 1; score for each taxon showing positive association with T2DM was defined as -1. We then summed up values from all taxa. We selected logistic regression model to estimate association of the MRS with T2DM risk, and linear model to estimate the association of the MRS with the continuous variables, adjusted for age, sex, dietary energy intake, alcohol intake and BMI at the time of sample collection.

### Clustering diseases

The clustering analysis was carried out with ‘cluster’ and ‘factoextra’ for plot on R. We performed PAM algorithm based on predicted abundance of diseases or average relative abundance after Z-score normalization ^83^. PAM algorithm searches k medoids among the observations and then found nearest medoids to minimize the dissimilarity among clusters ^18^. Given a set of objects *x* = (*x*_l_ *x*_2_, …, *x*_*n*_), the dissimilarity between objects *x_i_* and *x_j_* is denoted by d(*i,j*). The assignment of object i to the representative object j is denoted by *z_ij_*. *z_ij_* is a binary variable and is 1 if object i belongs to the cluster of the representative object j. The function to minimize the model is given by:

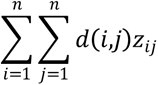

To identify the optimal cluster number, we used ‘pamk’ function in R to determine the optimum average silhouette width. For each object i, we defined *N_i_* as the average dissimilarity between object i and all other objects within its cluster. For the remaining clusters, b(*i,w*) represents the average dissimilarity between i and all objects in cluster *w.* The minimum dissimilarity *M_i_* can be calculated by:

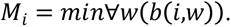

The silhouette width for object i can be calculated by:

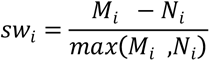

Then we calculated the average of silhouette width for each object. The cluster number is determined by the number of which the average silhouette width is maximum.

## Acknowledgments

This study was funded by National Natural Science Foundation of China (81903316, 81773416), Westlake University (101396021801) and the 5010 Program for Clinical Researches (2007032) of the Sun Yat-sen University (Guangzhou, China). The authors declare no conflict of interest. We thank the Westlake University Supercomputer Center for providing computing and data analysis service for the present project.

## Data availability

The raw data for 16 S rRNA gene sequences are available in the CNSA (https://db.cngb.org/cnsa/) of CNGBdb at accession number CNP0000829.

